# SequenceBouncer: A method to remove outlier entries from a multiple sequence alignment

**DOI:** 10.1101/2020.11.24.395459

**Authors:** Cory D. Dunn

**Affiliations:** Institute of Biotechnology, University of Helsinki, Helsinki, 00014, Finland

## Abstract

Phylogenetic analyses can take advantage of multiple sequence alignments as input. These alignments typically consist of homologous nucleic acid or protein sequences, and the inclusion of outlier or aberrant sequences can compromise downstream analyses. Here, I describe a program, SequenceBouncer, that uses the Shannon entropy values of alignment columns to identify and remove outlier entries in a manner responsive to overall alignment context. I demonstrate the utility of this software using alignments of mammalian reference mitochondrial genomes, bird cytochrome *c* oxidase-derived sequence barcodes, and COVID-19 sequences.

## Introduction

Outlier sequences within a dataset, which can include mis-annotated entries, mis-assembled sequences, and sequences containing rare structural rearrangements, can compromise our ability to perform robust phylogenetic analyses [1–3]. Avoidance of aberrant sequence inclusion within alignments becomes more challenging with the increasing rate of sequence acquisition.

To address this problem, I developed a method that could remove entire outlier sequences from a multiple sequence alignment in a manner responsive to overall sequence context and at moderate computational cost. This approach is incorporated into SequenceBouncer (https://github.com/corydunnlab/SequenceBouncer), a software which functions within a Python framework and takes advantage of Shannon entropy calculations to remove potentially undesirable alignment entries.

## Workflow and examples

### Pairwise sequence comparisons take advantage of alignment column Shannon entropy values to infer divergence

I first established a workflow in which all of the sequences within an input multiple sequence alignment are compared to one another in order to facilitate outlier identification. Hereafter, this workflow is designated a ‘full analysis’. As the initial step in this analysis, an input FASTA file is examined to determine the percentage of each alignment column that consists of gaps (Figure 1). Correct choice of a gap threshold (-g flag) by the user focuses downstream analysis upon generally well-aligned positions. A gap threshold that is excessively low leads to removal of information that might be helpful in identifying outlier sequences. A gap threshold that is too high will include alignment positions that may confound the analysis or squander computational resources on non-informative sites. The gap threshold default for SequenceBouncer is set at 2%.

**Figure 1:**
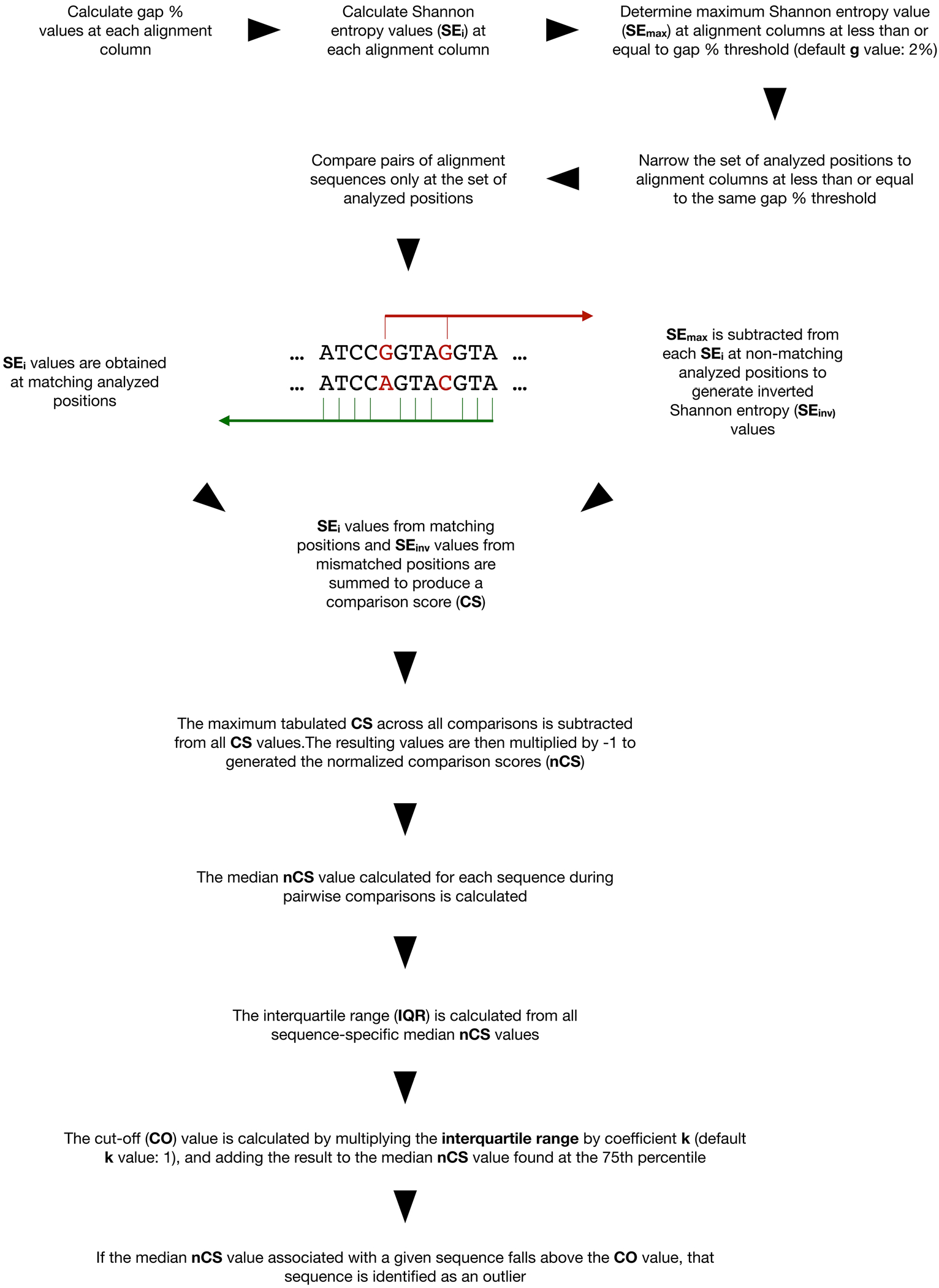
Alignment column Shannon entropy is deployed toward outlier sequence identification. A gap threshold for column analysis is selected, and the Shannon entropy for each alignment column (SE_i_) is calculated. All possible pairwise comparisons are performed, and where the sequences match, the SE_i_ values are added to the comparison score (CS). For mismatched positions, the maximum Shannon entropy (SE_max_) calculated across all positions below the gap threshold is subtracted from the column SE; value to generate ‘inverse’ Shannon entropy (SE_inv_) values, and these results are also added to the CS. Each CS is normalized (nCS), and the median nCS values across the comparisons to each sequence are used to determine a cut-off (CO) value above which a sequence is determined to be an outlier.

Next, the Shannon entropy value (SEi) associated with each alignment column, expressed in bits, is evaluated. Shannon entropy is linked to the amount of ‘surprise’ that one might encounter within a message with an expectation of ergodicity [4,5]. I use alignment columns here as ‘messages’ [6], and I take advantage of their Shannon entropy values as an appropriate basis for detecting unexpected divergence of a sequence from the bulk of alignment entries. The maximum SEi value obtained when considering all positions under the selected gap threshold (SE_max_) is saved for use in later calculations.

Following the calculation of SEi for each alignment column and determination of the SE_max_ value, pairwise comparisons of alignment sequences are initiated. If the SE_i_ value of an alignment column at which characters match is comparatively high, then the characters may be considered to be surprisingly identical, providing evidence for a greater level of similarity between the compared sequences. In contrast, if the SE_i_ value of a column at which characters match is low or zero, a character match is less noteworthy, given the high conservation at that position and the assumption that the user has attempted to align homologous sequences. The SE_i_ values obtained from matched positions are added together.

Next, positions divergent between the two compared sequences are considered. Here, a lack of a match at each position is penalized to a degree determined by the variability observed in that alignment column. Toward this goal, the SE_max_ value is subtracted from the SE_i_ value at each non-matching position to generate an ‘inverse’ (SE_inv_) score. SE_inv_ scores from non-matching positions are added to the sum of SE_i_ values obtained at matching positions to generate a total comparison score (CS). A higher CS score emerging from a pairwise comparison is linked to a more robust alignment and greater sequence relatedness.

Pairwise analyses of sequences continue, and CSs are generated, until each sequence within the alignment has been compared to every other sequence. The resulting CSs are then converted to normalized comparison scores (nCSs) linking higher value to greater divergence. To generate nCSs, the maximum CS obtained across all pairwise sequence comparisons is subtracted from all tabulated CSs, and then the resulting set of values is multiplied by −1.

Abnormally high nCS scores associated with a given entry across a majority of pairwise sequence comparisons signals outlier status. A cut-off (CO) value is calculated from the nCS dataset in order to discriminate alignment outliers. To ascertain this CO value, the median nCS obtained for each sequence across all of its pairwise comparisons is determined, and the interquartile range is calculated across these median nCSs. The interquartile range is then multiplied by the coefficient k (-k flag), which can either be supplied by the user or left at the default value of 1. The result is then added to the nCS found at the 75th percentile to produce the CO score. Those sequences for which median nCSs obtained across its comparisons with other sequences lies above the CO value ‘fail’ testing by SequenceBouncer and will be marked for potential removal from the alignment.

When the is completed, results are reported within a comma-separated values (CSV) file that lists each input sequence accession and whether the median nCS for that sequence across pairwise comparisons is above the outlier threshold. In addition, a ‘cleansed’ FASTA file lacking outlier sequences, as well as a FASTA file containing rejected outlier sequences, are provided as output. Finally, when a full analysis is performed, a complete table of nCSs emerging from all pairwise comparisons is supplied to the user.

### Example 1 Mammalian mitochondrial DNA sequences

Sequences found within the RefSeq database [7] are subject to validation and curation. However, it is clear that some organellar sequences available within this database contain errors, and others harbor rearrangements, insertions, and duplications that would require further treatment before a serviceable alignment can be generated [8]. Accordingly, I expected that the set of mammalian mitochondrial DNA sequences retrieved from RefSeq might provide a useful test dataset when developing my approach to outlier sequence detection.

After downloading mammalian mtDNA sequences obtained from RefSeq and removing duplicate sequences, I aligned the resulting dataset using MAFFT [9]. Next, I performed a full analysis using SequenceBouncer to detect and remove outliers. Positions gapped at 2% of sequences or less were analyzed, and nCS calculations for all possible pairwise sequence comparisons were completed (Figure 2A). When plotting the median nCS associated with each sequence (Figure 2B) a subset of entries was clearly associated with excessive median nCSs (nCS > 8500), and visual inspection of these sequences indeed indicated poor alignment with the vast majority of other sequences. A few of these sequences appear to be inverted with respect to the majority of other downloaded RefSeq mammalian mtDNA entries. I also noted that the 37 marsupial mtDNAs and three monotreme mtDNAs included in this alignment of 1303 mammalian mtDNAs were associated with median nCSs that were elevated relative to the bulk of entries (7800 < nCS < 8500), as might be expected when considering a sequence set heavily biased toward placental mammals. Input files and SequenceBouncer output related to the examples provided here can be found at (doi: 10.5281/zenodo.4285789). This analysis was performed using a MacBook Pro with a 2.8 GHz Quad-Core Intel Core i7 processor and 16 GB of 1600 MHz DDR memory, and the total time required to complete this run of SequenceBouncer using the mammalian mtDNA input alignment was ~ 5 minutes.

**Figure 2:**
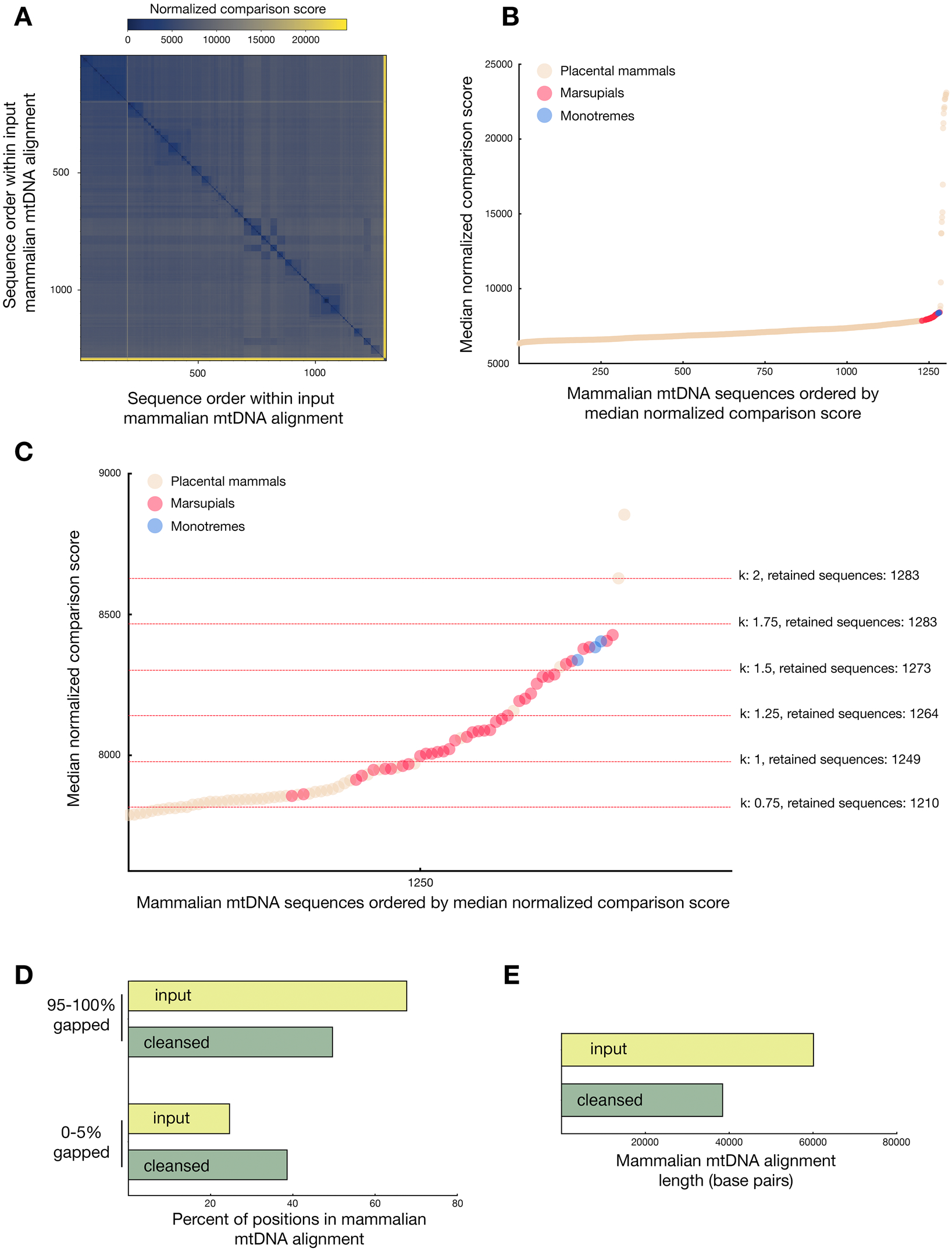
Identification of outlier sequences found within an alignment of mammalian mtDNA sequences. SequenceBouncer was run on an input alignment of mammalian mtDNA sequences (1303 entries) using a gap threshold of 2%, and all pairwise sequence comparisons were performed. (**A**) SequenceBouncer output showing the nCS associated with each pairwise sequence comparison. Blue reflects higher identity, and yellow signals higher divergence. (**B**) A subset of input mtDNA sequences are clearly outliers from the alignment. Mammalian mtDNA sequences are ordered by increasing median nCSs, and infraclasses are reflected by dot color. (**C**) k settings shift the CO value for outlier detection. The CO value obtained using the mammalian mtDNA dataset is indicated at k values ranging from 0.75 to 2 in steps of 0.25. (**D**) Removal of outlier sequences leads to a decrease in heavily gapped columns and an increase in minimally gapped columns upon realignment. The cleansed output alignment obtained at a k value of 1.75 (1283 sequences) was realigned, and the gap frequency was calculated for input alignment and cleansed, re-aligned output. (**E**) Removal of outlier sequences allows for a reduction in alignment sequence length. SequenceBouncer analysis was performed as in (**D**), and the length of cleansed, realigned entries was assessed.

Next, I explored the outcome of changing k, the coefficient used to calculate CO values. At a k value of 0.75, the most poorly aligned sequences were removed, as well as all monotreme and marsupial sequences (Figure 2C). In contrast, at k values of 1.75 and above, all monotreme and marsupial sequences were retained, and only clearly aberrant sequences were removed. These tests demonstrate that SequenceBouncer output can be easily tailored, by use of the k variable, to the user’s desired definition of ‘outlier’.

I used simple metrics to investigate whether the output sequence set might have been improved by SequenceBouncer analysis. When aberrant sequences are included in an alignment, the proportion of highly gapped alignment columns is often elevated due to the accommodation of these outlier entries [10]. Consequently, removal of outlier sequences should lead, upon a repeat of the alignment process, to a more compact alignment with a reduced number of highly gapped alignment columns and a lower number of total columns. When comparing my initial input alignment to the re-aligned output of SequenceBouncer run using a k setting of 1.75, many abundantly gapped columns were removed (Figure 2D), and the overall length of the aligned sequences was reduced (Figure 2E), indicating the successful generation of an output dataset harboring entries with greater overall sequence similarity.

### Analysis of large alignments by sequence sampling

When alignment depth and length are moderate, as in the case of the mammalian mtDNA sequence set, all sequences can be compared to all other sequences within minutes by use of a laptop or desktop computer. However, more expansive alignments, which may encompass tens of thousands of sequences containing tens of thousands of characters, begin to tax the computational resources and patience available to researchers who would like to quickly and effectively remove non-homologous entries. Consequently, I incorporated into SequenceBouncer the possibility of ‘sampling-based’ approaches that permit outlier detection, yet do not require an extraordinary amount of computational effort.

Here, within each ‘trial’, samples are selected from the alignment (Figure 3A), and the size of each sample is chosen by the user (-n flag). Whether the median nCS for a sequence is higher than the CO value is calculated as if the chosen sample represented the entire input alignment. Sequences are sampled without replacement to the starting dataset until no more sequences remain to be selected and analyzed from the larger, source alignment.

**Figure 3:**
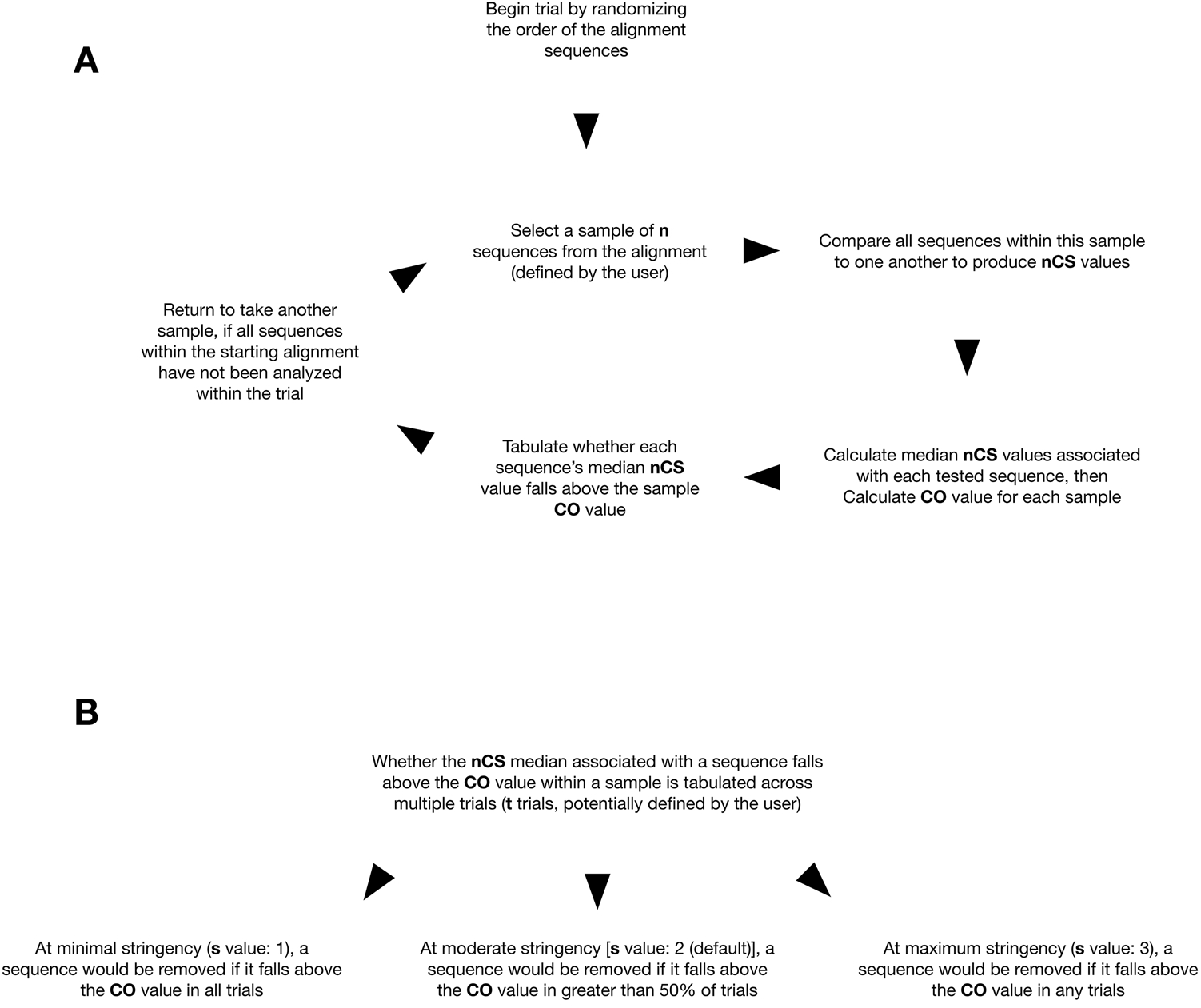
A sampling-based analysis can permit high prediction precision at reduced computational cost. (**A**) A subset of sequences, with the sample size determined by the user, is selected from the full alignment, and whether the nCS falls above the CO score is determined as in Figure 1. (**B**) Multiple trials, with the number determined by the user, can be performed to increase the certainty of outlier selection. Stringency, related to the number of trials during which a sequence fell above the CO value, can be set based upon the goals of the user.

A single trial of each sequence is unlikely to be sufficient for confident identification of outliers under most use cases of this sampling-based approach. Therefore, the user may request multiple trials (-t flag). The outcomes of these trials, along with the stringency (-s flag) setting, are used to determine whether a sequence should be removed from the alignment. When SequenceBouncer is set to minimal stringency (-s 1), then a sequence must fail all trials to be removed from the alignment. When the software is set to moderate stringency (-s 2), a sequence must fall above the CO value in more than 50% of the trials in order to be selected for removal from the alignment. At maximum stringency (-s 3), a sequence is excluded from the cleansed alignment if it is flagged as an outlier in any trials. When a sampling-based approach is applied, the output CSV file lists accessions, the number and fraction of trials during which a sequence was identified as an outlier, and whether, based upon the stringency setting, a sequence was retained in a cleansed FASTA output alignment. Random shuffling of sequences in the source alignment occurs before samples are extracted, and a random seed can be set by the user (-r flag) to allow reproducibility between runs.

Below, I demonstrate how the SequenceBouncer program can take advantage of a sampling-based approach to remove outliers from larger alignments.

### Example 2 Bird cytochrome c oxidase-derived barcodes

A portion of the well-conserved gene encoding cytochrome *c* oxidase subunit I is often deployed as a sequence barcode during ecological studies [11–13]. These barcodes, as well as details of sample acquisition, are readily available from the Barcode of Life Data (BOLD) Systems database [14]. However, not all of the barcode sequences found within this database are properly annotated and of high quality. I tested whether SequenceBouncer might successfully remove outlier barcodes, and I specifically focused upon bird COI-5P entries, which I found to be of utility during a recent study seeking hummingbird-specific changes to mtDNA-encoded proteins [15].

Toward this goal, I downloaded barcode data for class Aves from the BOLD database, deleted duplicate barcodes and barcodes not annotated as being of the COI-5P class, then aligned the remaining sequences using MAFFT [9]. Next, I performed a full analysis that carried out all pairwise sequence comparisons (Figure 4A). The gap threshold was set at 5%, and the k coefficient was set to 1. This full analysis required nearly 5 hours on the above-mentioned test system. Upon re-alignment of the cleansed dataset, the fraction of highly gapped columns (95-100% gapped) was reduced, and the fraction of seldom gapped columns (0-5% gapped) was increased (Figure 4B), indicating successful removal of sequences likely to be aberrant. Moreover, the length of the sequences within the cleansed and realigned dataset was reduced (Figure 4C), also suggesting removal of outliers potentially compromising the initial alignment. Finally, the distribution of the ungapped sequence lengths of entries within the cleansed alignment clearly demonstrated removal of short sequences of a length inconsistent with the length of a useful COI-5P barcode (Figure 4D).

**Figure 4:**
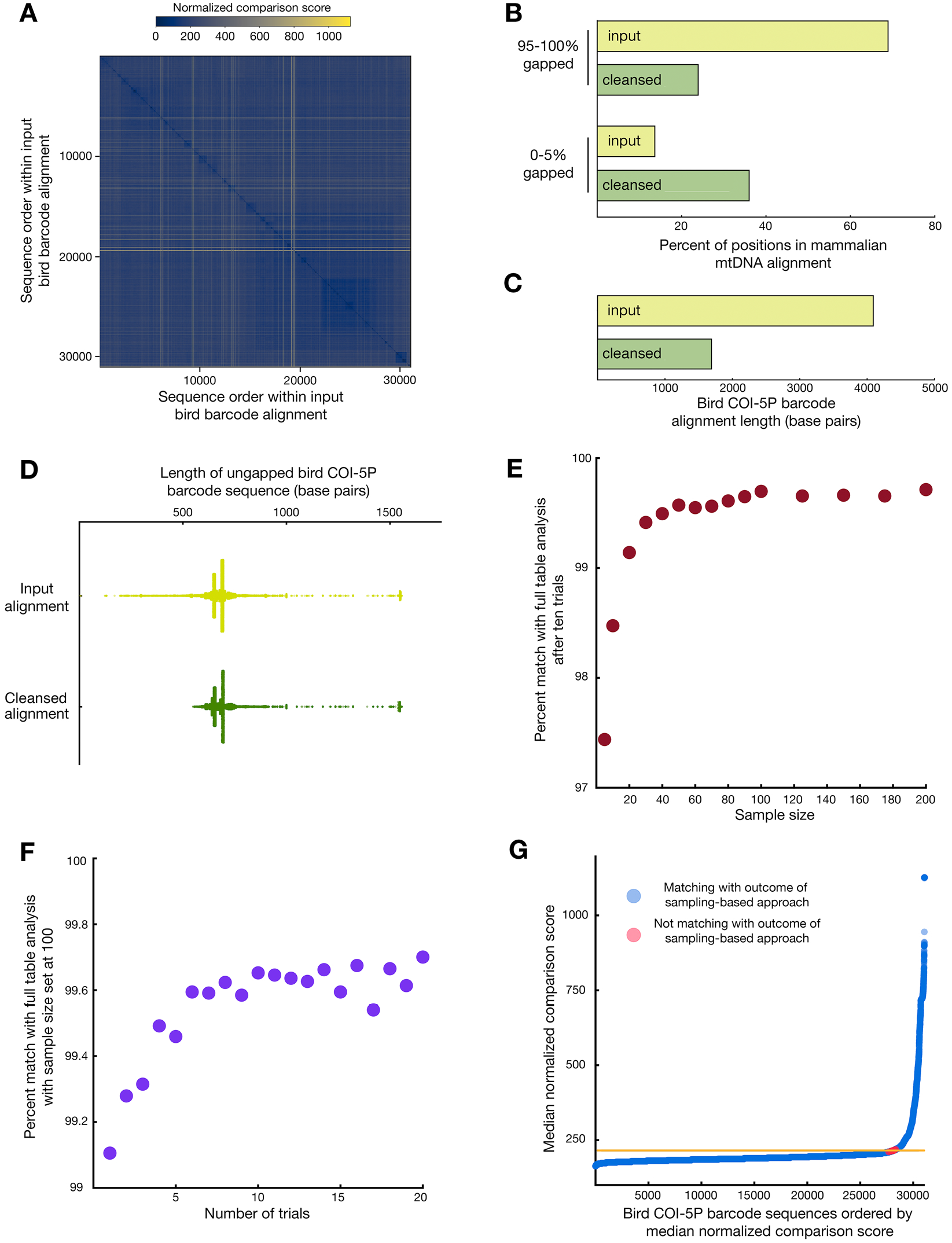
Identification of outlier sequences within an alignment of bird COi-SP barcodes. (**A**) An alignment of bird COl-5P barcodes (31,087 sequences) was analyzed with SequenceBouncer at a gap threshold of 5% and k setting of 1. A full analysis consisting of all pairwise sequence comparisons was performed. The graphical output file is provided, with blue linked to higher identity between two sequences and yellow signaling higher divergence. The cleansed output alignment (28,141 sequences) and the input alignment were realigned and processed (**B**) as in Figure 2D or (**C**) as in Figure 2E, as fewer highly gapped alignment columns and shorter aligned sequences can indicate improved homology between alignment entries. (**D**) Short sequences unlikely to encode a full-length COl-5P barcode are depleted from the input alignment using SequenceBouncer. Input and cleansed output alignments were de-gapped, and the distribution of resulting sequence lengths is shown. (**E**) An exploration of sample size during a sampling-based approach. The alignment of bird COl-5P barcodes was used as input for SequenceBouncer with k setting of 1, a gap threshold of 5%, a stringency setting of ‘moderate’, and a trial number of 10. The percent of outlier determinations from the sampling-based approach that match with the output of a full analysis that takes advantage of all pairwise sequence comparisons is shown. (**F**) An exploration of trial number during a sampling-based approach. The alignment of bird COl-5P barcodes was used as input with k setting of 1, a gap threshold of 5%, a stringency setting of ‘moderate’, and a sample size of 100. The percent of outlier determinations matching with the output of a full analysis after the indicated number of trials is reported. (**G**) Entries for which the outcome of a sampling-based analysis does not match the full analysis lie near the CO value found during the full analysis. Bird COl-5P barcodes are ordered by increasing median nCSs. Samples for which the outcome of a full analysis matches or does not match the sampling-based decision on sequence retention are labelled in blue and red, respectively. The CO value obtained during a full alignment analysis with all pairwise sequence comparisons is designated by an orange line.

Because an analysis taking advantage of all pairwise sequence comparisons was computationally costly, I explored whether my sampling-based approach could also successfully identify bird barcode outliers, but at a fraction of computational effort. First, I tested which sample size, given the input alignment and 10 trials at moderate stringency, would permit a close approximation of results obtained when performing the full analysis of all pairwise sequence comparisons. I found that the percent of outlier calls that matched during comparison of sampling-based tests with the full analysis at a sample size of ~ 100 sequences appeared to plateau at ~ 99.7% (Figure 4E). Of course, results achieved using any single input alignment will not necessarily generalize to every possible dataset, and so each SequenceBouncer user may assess the stability of results obtained on their own input dataset at different parameter settings.

Next, I examined correspondence between sampling-based and full analyses as the number of trials was varied. Gap threshold and k values were maintained at 5% and 1, respectively. The sample size was set to 100, stringency was set to ‘moderate’, and trial numbers between 1 and 20 were tested. A greater number of sampling-based trials was clearly associated with improved correspondence to the full pairwise analysis. A plateau of ~ 99.7% correspondence between these two analyses was obtained at a trial number of ~ 10 (Figure 4F). Again, the most appropriate parameters would depend upon the specific input alignment, yet an increase in trial number can clearly lead to improved results. As expected, those entries for which the sampling-based and full analysis output did not match were sequences for which the median nCS across all pairwise comparisons was close to the full analysis CO value (Figure 4G). Excitingly, the time to complete ten trials of the input bird COI-5P alignment at a sample size of 100 and a gap threshold of 5% was only ~ 10 minutes on the test system, a greater than 20-fold decrease in required computational time when compared to a full analysis.

### Example 3 COVID-19 sequences

The examples provided above demonstrate that mammalian full-length mtDNA sequence alignments of moderate depth, as well as high-depth alignments consisting of short barcodes, can be effectively and rapidly processed by SequenceBouncer. Next, I turned my attention to a rapidly growing set of COVID-19 sequences available for download from the RefSeq database. The COVID-19 RNA is approximately 30,000 bases in length, and my input alignment consisted of nearly 37,500 entries, most of which are nearly identical or identical to other sequences within the dataset [16]. It is common to find sequencing errors in COVID-19 sequences that might compromise phylogenetic analyses [17], and I used a sampling-based approach to determine which COVID-19 sequences should be marked as outliers and removed from downstream analysis.

While analyzing these COVID-19 sequences, I explored the outcomes of different SequenceBouncer stringency settings. Gap threshold was set to 2%, k value was set to 1, sample size was set to 100, and trial number was set to 10. As discussed above, the stringency setting directs exclusion of entries from output alignments based upon the proportion of trials for which an entry is determined to be an outlier. At minimal stringency, a substantial number of truncated sequences were removed from the alignment (Figure 5A and Figure 5B). The range of ungapped sequence lengths was, as expected, further narrowed by performing the analysis of COVID-19 sequences as moderate and maximum stringency. The COVID-19 alignment mostly consists of very similar sequences, and so comparatively few columns are almost totally gapped, in contrast to the mammalian mtDNA and bird barcode alignments. Nonetheless, realignment after SequenceBouncer processing clearly led to a reduction in gapped alignment positions that accommodate outlier sequences (Figure 5C), particularly at moderate and maximal stringency settings. Of note, a set of corrupted COVID-19 sequences, LR897977.1 - LR898047.1 (https://mafft.cbrc.jp/alignment/software/) were identified as outliers when SequenceBouncer was run at moderate and maximum stringency, further supporting the utility of my approach. Improvements in alignment quality as stringency was progressively increased are also clear when alignments are graphically represented (Figure 6). On the test system, analyses took only ~ 120 minutes. Clearly, SequenceBouncer output can remove aberrant entries from very sizable datasets of highly homologous, virus-derived sequences.

**Figure 5:**
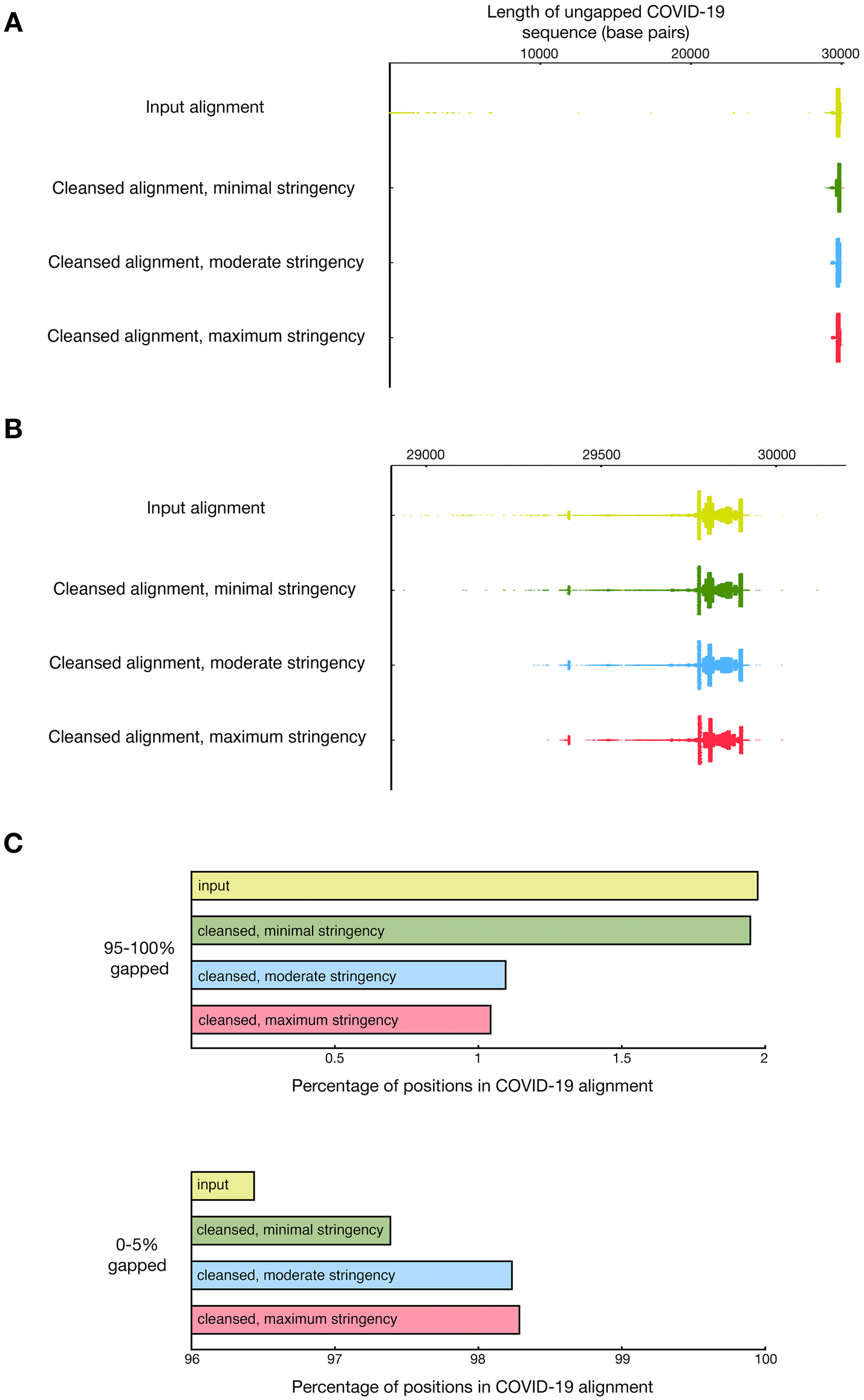
Identification of outlier sequences within an alignment of COVID-19 sequences. SequenceBouncer was used to analyze an input alignment by a sampling-based approach over 10 trials at a gap threshold of 2%, a k setting of 1, and a sample size of 100. The analyses were performed at minimal, moderate, and maximum stringency. (**A**) Apparently truncated sequences shorter than the reference sequence are removed from the input alignment (37,470 sequences) at minimal stringency (33,351 sequences remain), moderate stringency (29,821 sequences remain), or maximum stringency (27,147 sequences remain). (**B**) as in (**A**), but showing a range with minimum and maximum approaching the COVID-19 reference sequence length. (**C**) Removal of COVID-19 outlier sequences leads to a decrease in heavily gapped columns and an increase in minimally gapped columns upon realignment. cleansed alignments were realigned, and the fraction of heavily gapped columns (95-100% of samples) and seldom-gapped columns (0-5% of samples) was calculated.

**Figure 6:**
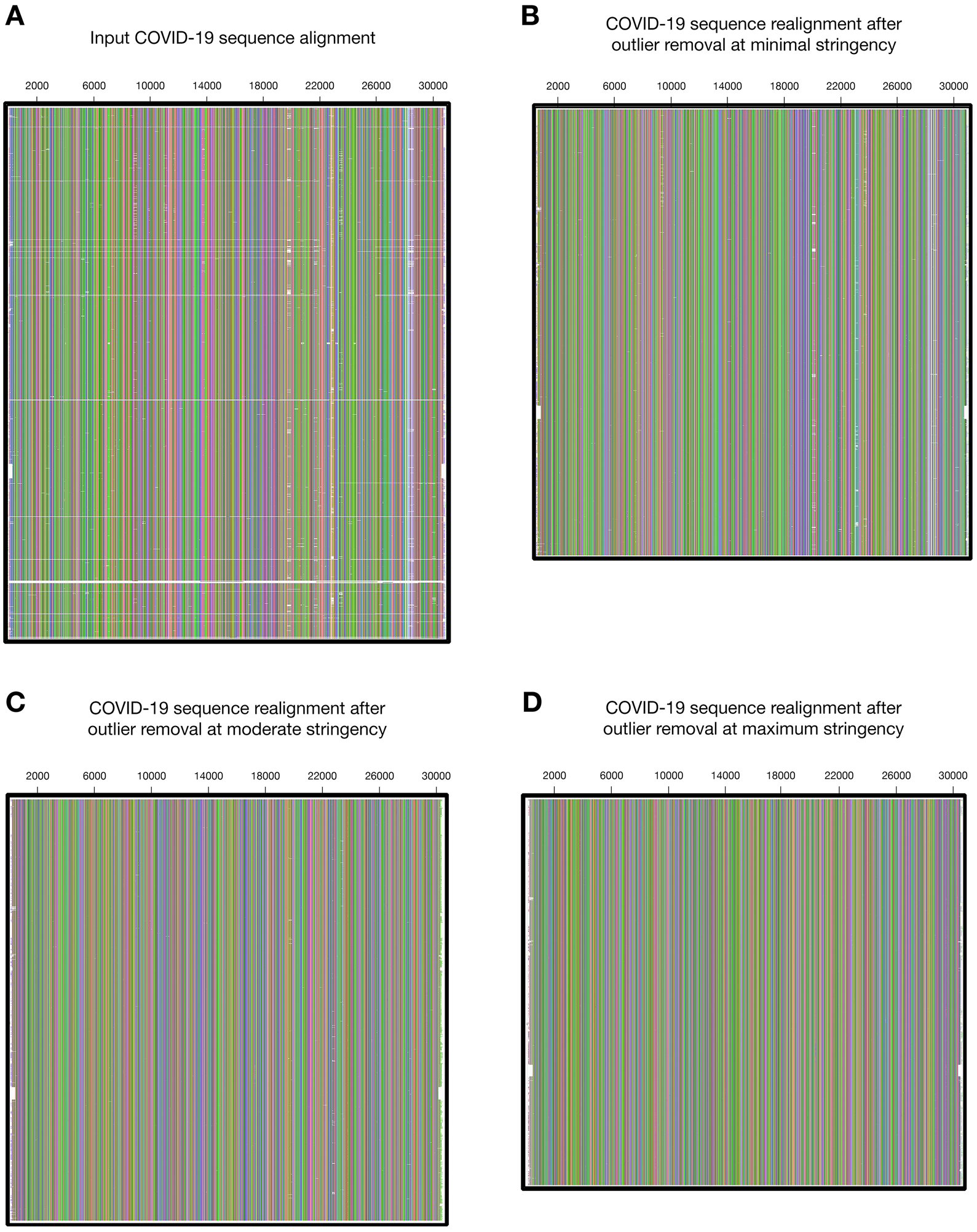
COVID-19 alignment quality appears to be augmented as SequenceBouncer stringency is increased. The COVID-19 input alignment (**A**), as well as the realignments of files cleansed by a sampling-based methodology at minimal (**B**), moderate (**C**), and maximum (**D**) stringency are visualized using AliView (31] at a comparable level of magnification.

## Conclusion

I report the development of SequenceBouncer, which detects and removes outlier entries from a multiple sequence alignment. Our approach joins other methods that may allow detection of potentially erroneous entries or selected problematic regions in multiple sequence alignments or among COI barcodes [18–24].

SequenceBouncer takes advantage of Shannon entropy computations to allow a context-dependent calculation of which sequences might not be homologous with other entries. Some outliers are mis-annotated, mis-assembled, contain many sequencing errors, or are not oriented appropriately when considering the greater majority of other analyzed sequences. Other outliers might contain important structural rearrangements not found within the bulk of other alignment entries [25].

This software, written in Python 3, is uncomplicated to install and run, requiring only Numpy and Pandas as dependencies. Moreover, key parameters can be tailored to the specific needs of the user. Depending upon the input alignment size, analyses either can be performed by completion of all pairwise sequence comparisons or, if computational resources are limiting, can be executed by a rapid and reliable sampling-based approach. Even a large COVID-19 input alignment of 37,470 sequences, each consisting of 31,012 nucleotide positions, could be effectively processed within a few hours using a laptop computer.

## Methodology

### Sequence acquisition

Mammalian mtDNA RefSeq accessions (RefSeq release 202) were downloaded on September 29, 2020 from the Organelle Genome Resources page of NCBI (https://www.ncbi.nlm.nih.gov/genome/organelle/), and the 1316 associated full length mtDNA sequences were obtained by use of the NCBI Entrez utilities (https://www.ncbi.nlm.nih.gov/books/NBK25501/). Duplicate sequences were removed by use of seqkit v0.13.2 [26] to generate a file containing 1303 mammalian mtDNA entries.

Bird COI-5P barcodes were downloaded from the BOLD database [14] on October 21, 2020 using the query ‘Aves’. 46,397 entries were filtered for those containing COI-5P in their description line, then duplicate sequences were deleted by seqkit v0.13.2 [26] to generate a file for initial alignment and subsequent SequenceBouncer testing that contained 31,087 barcodes.

Full length COVID-19 sequences were downloaded from the severe acute respiratory syndrome coronavirus 2 data hub (https://www.ncbi.nlm.nih.gov/labs/virus/vssi/#/virus?SeqType_s=Nucleotide&VirusLineage_ss=SARS-CoV-2,%20taxid:2697049) on November 9, 2020. Here, duplicate sequences were retained during downstream analysis.

### Sequence alignment

For full-length mammalian mtDNA sequences and bird COI-5P barcodes, initial alignment and alignment after SequenceBouncer processing were carried out by MAFFT v7.471 [9] using the FFT-NS-2 approach. For bird COI-5P barcodes, the ‘anysymbol’ flag was activated.

For full-length COVID-19 sequences, initial alignment and realignment following SequenceBouncer processing was performed using an online MAFFT server tailored to the analysis of datasets consisting of viral sequences with a high level of identity (https://mafft.cbrc.jp/alignment/software/closelyrelatedviralgenomes.html) [27]. Sequences with greater than 5% ambiguous letters were removed, a ‘6merpair’ strategy was deployed, ‘anysymbol’ and ‘memsave’ modes were activated, and the scoring matrix selected was ‘1PAM/kappa=2’. The COVID-19 reference sequence, NC_045512.2, was used as the alignment to which all other sequences were added.

### Gap quantification and removal

De-gapping of sequences and sequence length calculations was performed by use of seqkit v0.14.0 [26]. Calculation of alignment column gap frequencies for analysis of SequenceBouncer results was carried out using trimal v1.4.rev15 [28].

### Taxonomic analysis

Taxonomy analysis was performed using the’taxize’ package [29] with the NCBI taxonomy database [30].

### Data visualization

Graphs were produced by use of Prism 9.0.0 (https://www.graphpad.com), and COVID-19 alignment visualization was achieved using AliView 1.24 [31].

## Acknowledgements

This work was funded by a Senior Researcher grant from the Sigrid Jusélius Foundation to C.D.D. I thank Gülayşe İnce Dunn, Svetlana Konovalova, Tamara Somborac, and Péter Poczai for helpful comments on the manuscript. I also thank Dave Whipp and Vuokko Heikinheimo for allowing me to attend their Geo-Python course.

